# Coordinated conformational changes in the V_1_ complex during V-ATPase reversible dissociation

**DOI:** 10.1101/2021.11.09.467972

**Authors:** Thamiya Vasanthakumar, Kristine A. Keon, Stephanie A. Bueler, Michael C. Jaskolka, John L. Rubinstein

**Affiliations:** Molecular Medicine Program, The Hospital for Sick Children; Department of Biochemistry, The University of Toronto; Department of Medical Biophysics, The University of Toronto; Department of Biochemistry and Molecular Biology, SUNY Upstate Medical University, Syracuse, NY 13210

## Abstract

Vacuolar-type ATPases (V-ATPases) are rotary enzymes that acidify intracellular compartments in eukaryotic cells. These multi-subunit complexes consist of a cytoplasmic V_1_ region that hydrolyzes ATP and a membrane-embedded V_O_ region that transports protons. V-ATPase activity is regulated by reversible dissociation of the two regions, with the isolated V_1_ and V_O_ complexes becoming autoinhibited upon disassembly and subunit C subsequently detaching from V_1_. In yeast, assembly of the V_1_ and V_O_ regions is mediated by the RAVE complex through an unknown mechanism. We used cryoEM of yeast V-ATPase to determine structures of the intact enzyme, the dissociated but complete V_1_ complex, and the V_1_ complex lacking subunit C. Upon separation, V_1_ undergoes a dramatic conformational rearrangement, with its rotational state becoming incompatible for reassembly with V_O_. Loss of subunit C allows V_1_ to match the rotational state of V_O_, suggesting how RAVE could reassemble V_1_ and V_O_ by recruiting subunit C.

## Introduction

Vacuolar-type ATPases (V-ATPases) are ATP-hydrolysis-driven proton pumps that acidify intracellular compartments in most eukaryotic cells^1^. This activity makes V-ATPases essential for fundamental cellular processes including protein biosynthesis and post-translational modification, intracellular trafficking, endocytosis and exocytosis, and lysosomal degradation^1,2^. V-ATPases can also localize to the plasma membrane of specialized cells, such as osteoclasts^3^ and intercalated cells of the kidney^4^, where they acidify the extracellular environment. Aberrant expression or activity of the enzyme has been associated with cancer metastasis^5^ and neurodegeneration^6^, and loss-of-function mutations in specific subunit isoforms can cause genetic diseases such as osteopetrosis^7^ and distal renal tubular acidosis^8^.

The multi-subunit V-ATPase consists of a soluble V_1_ region and a membrane-embedded V_O_ region (Fig. 1A, *left*). Three pairs of catalytic A and B subunits in the V_1_ region hydrolyze ATP, leading to conformational changes that induce rotation of a rotor subcomplex (subunits DFdc_8_c′c″Voa1p) that extends into the lipid bilayer to form part of the V_O_ region^9–11^. Three peripheral stalks, consisting of heterodimers of subunits E and G, link the catalytic A_3_B_3_ hexamer to a collar structure formed from subunits C, H, and the soluble N-terminal domain of subunit a. These peripheral stalks prevent subunits a, e, and f in the V_O_ region from rotating along with the c_8_c′c″-ring (c-ring). Rotation of the c-ring against the membrane-embedded C-terminal domain of subunit a drives proton translocation through V_O_. Protons enter V_O_ through a cytosolic half-channel between subunit a and the c-ring, sequentially neutralize conserved glutamate residues on each subunit of the rotating c-ring, and exit V_O_ through a luminal half-channel in subunit a^10^. In this way, ATP hydrolysis in the V_1_ region is coupled to proton transport through the V_O_ region. CryoEM has shown that even in the absence of ATP the enzyme adopts three main conformations, corresponding to rotational positions of the rotor and known as rotational ‘State 1’, ‘State 2’, and ‘State 3’^9,12^.

**Fig 1.**
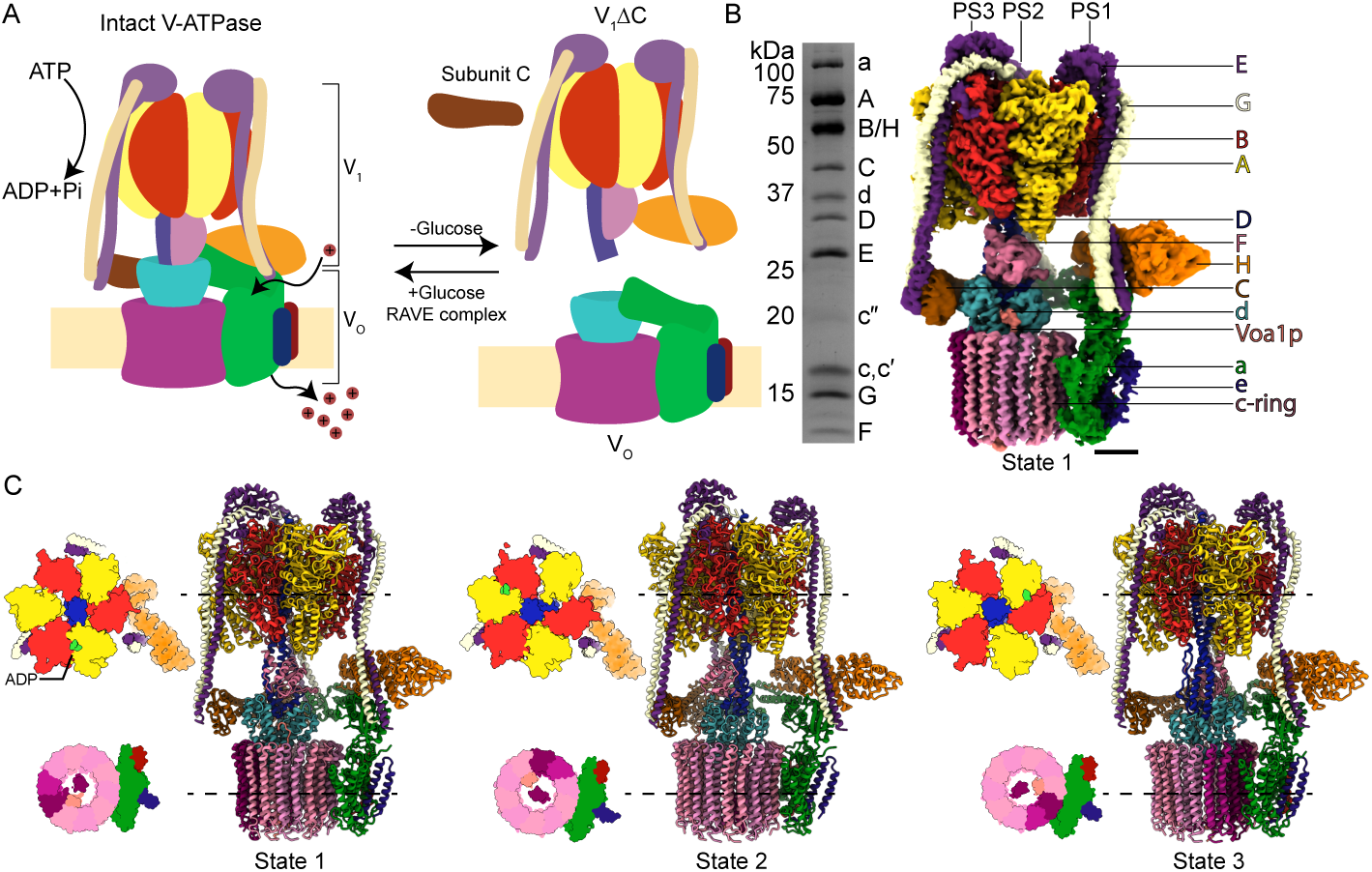
Structure of intact yeast V-ATPase. **A**, Glucose starvation induces dissociation of intact V-ATPase (*left*) into the V_1_ and V_O_ complexes with subunit C subsequently detaching (*right*). The RAVE complex can reassemble the complex. **B**, SDS-PAGE of the purified intact V-ATPase (*left*) allows high-resolution structure determination by cryoEM (*right*). A composite map is shown from local refinement of V-ATPase in rotational State 1. Scale bar, 25 Å. **C**, Atomic models for V-ATPase in rotational State 1 (*left*), State 2 (*middle*), and State 3 (*right*) with cross-sections through the V_1_ region and V_O_ regions shown for each state. An ADP molecule (*green*) is seen in each V_1_ region.

V-ATPase activity is controlled by regulated separation and reassembly of the V_1_ and V_O_ regions^13,14^. In yeast, V-ATPase dissociation was observed in response to glucose starvation, with restoration of glucose in the growth medium allowing reassembly^13^ (Fig. 1A, *right*). V-ATPase assembly in yeast is also influenced by osmotic stress^15^ and changes in extracellular pH^16^, while in mammalian cells regulated dissociation or assembly have been observed in response to changes in glucose and amino acid concentration^17,18^ or cell maturation^19^. Subunit C is released from the V_1_ region following separation from V_O13_ but co-purifies with isolated V_1_ with variable stoichiometry^20–22^. Once separated, the V_1_ and V_O_ complexes become autoinhibited and no longer hydrolyze ATP or transport protons, respectively^23,24^. Silencing of ATP hydrolysis in V_1_ requires subunit H^23^, which possesses N- and C-terminal domains (H_NT_ and H_CT_) connected by a flexible linker^25,26^. A crystal structure of the yeast V_1_ complex lacking subunit C (V_1_ΔC) suggested that H_CT_ undergoes a 150° rotation upon dissociation, contacting one of the catalytic AB pairs to inhibit ATP hydrolysis^27^.

The V_1_ΔC crystal structure shows the complex in rotational State 2^27^, whereas cryoEM of the dissociated V_O_ region shows it in State 3^10^. Consistent with this mismatch, the V_1_ and V_O_ regions do not spontaneously assemble *in vitro*, but assembly can be induced by replacing yeast subunit H with a chimeric protein consisting of the N-terminal domain of yeast subunit H and the C-terminal domain of human subunit H^28^. Both the initial assembly of V_1_ and V_O_ upon synthesis and reassembly following separation require the RAVE complex, composed of Rav1p, Rav2p, and Skp1p in yeast^29,30^. Rav1p is homologous with mammalian and insect rabconnectin-3α, which is involved in endosomal acidification and V-ATPase assembly^31,32^. RAVE binds free V_1_ complexes and associates with vacuolar membranes in a glucose-dependent manner^33^. Rav1p interacts with subunits E, G, and C of the V_1_ complex, as well as the soluble N-terminal domain of the vacuolar isoform of subunit a (Vph1p) from V_O_, and may promote assembly by bringing V_1_, V_O_, and subunit C together^33^. *In vitro*, RAVE accelerates reassembly of V_O_ with V_1_ containing chimeric subunit H^34^.

We used cryoEM to determine structures for the intact V-ATPase, a V_1_ complex including subunit C (‘complete’ V_1_), and a V_1_ complex lacking subunit C (V_1_ΔC). The complete V_1_ complex is found in State 2 and adopts a conformation that differs dramatically from the V_1_ region of intact V-ATPase, with both subunits C and H collapsed toward the central rotor. The conformational changes in subunits C and H, as well as adoption of rotational State 2, make the complete V_1_ complex incompatible for reassembly with V_O_ in State 3. In contrast, we find that the V_1_ΔC complex can adopt State 1, 2 or 3, with most complexes in State 3. V_1_ΔC in State 3 remains incapable of reassembling with V_O_ in State 3 due to the lack of subunit C. In support of this hypothesis, purification of V_1_ΔC from a yeast strain with the gene for subunit C deleted leads to copurification of Rav1p, Rav2p, and Skp1p, suggesting a frustrated RAVE-V_1_ΔC complex unable to assemble with V_O_ until subunit C is recruited. Together, these structures reveal dramatic conformational changes throughout V_1_ upon separation from V_O_ and suggest the sequence of events that occurs during the regulated reversible dissociation of V-ATPases.

## Results

### High-resolution structures of intact V-ATPase enable analysis of V_1_ conformational changes

While high-resolution structures of mammalian V-ATPase have been determined^12,35,36^, high-resolution structures of the complete yeast enzyme are not available in the protein data bank. Therefore, to facilitate comparison of the structure of the V_1_ region within intact V-ATPase with the structure of V_1_ when dissociated from V_O_, we used a previously-described protein purification^25^ with recent imaging and image analysis methods^12,37,38^ to determine high-resolution structures of the three main rotational states of intact yeast V-ATPase (Fig. 1B, Fig. S1A and B, Table 1). The resulting 3D maps had nominal overall resolutions of 3.5 Å for State 1 and 3.8 Å for State 2 and 3, with local resolutions ranging from approximately 3.1 to 6.8 Å for State 1, 3.3 to 7.1 Å for State 2, and 3.4 to 8.3 Å for State 3 (Fig. S1C). From the final dataset of intact V-ATPase particle images, 51% contributed to State 1, 28% to State 2, and 21% to State 3, consistent with previous studies^9,12^. Local resolutions were improved with focused refinement for different regions of the complex (Fig. S1D). Composite maps from combining focused refinements were then used to build atomic models for the three rotational states (Fig. 1C, Table 1), including modelling of an adenosine diphosphate (ADP) molecule in one catalytic site of each map (Fig. 1C, *light green*).

**Table 1.**
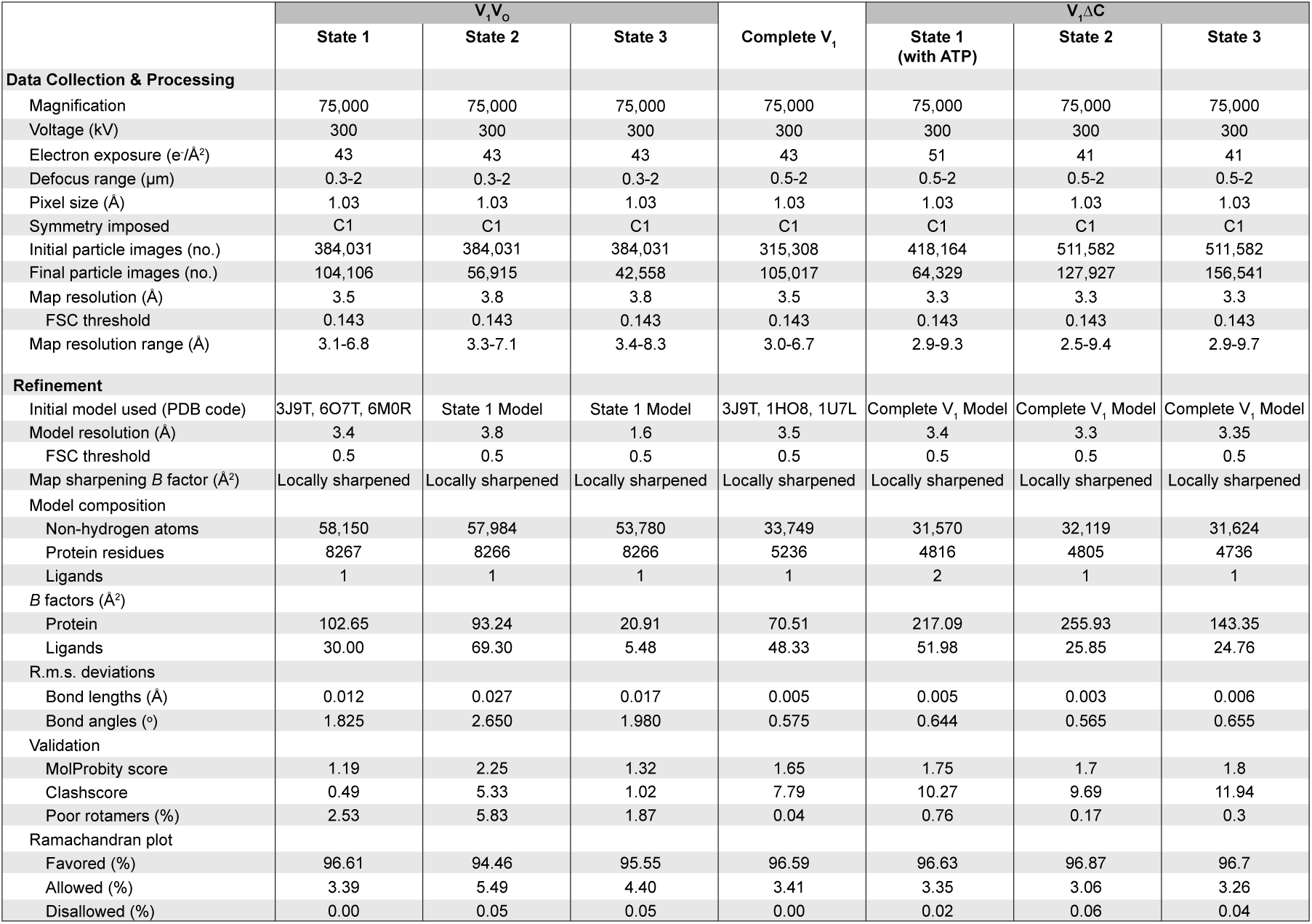
CryoEM and atomic model building statistics.

### Complete V_1_ including subunit C changes conformation upon separation from V_O_

To purify the V_1_ complex, we grew yeast to saturation to deplete glucose and induce dissociation of the V_1_ and V_O_ regions. Although subunit C is known to dissociate from V_1_ ^13^, it can still be found in many preparations of the complex^20–22^ including the one obtained here (Fig. 2A, *red*). The protein preparation was subjected to cryoEM (Fig. S2A and B, Table 1) and 3D classification was used to separate images of V_1_ complexes with and without subunit C, which we refer to as ‘complete V_1_’ and ‘V_1_ΔC’, respectively. This procedure resulted in a map of the complete V_1_ complex at a nominal overall resolution of 3.5 Å, with local resolutions ranging from 3.2 to 7.0 Å (Fig. 2B, Fig. S2C). The best resolved regions of the map were in the catalytic core and the worst resolution was in the collar region that contains subunit H and C, suggesting that the collar region of the dissociated V_1_ complex is flexible. Three-dimensional variability analysis^39^ also indicated movement in this region of the complex (Supplementary Video 1). The map of the complete V_1_ complex enabled de novo construction of atomic models for most of the subunits, with crystal structures for the subunits in the collar region flexibly fit into the map (Fig. 2C, Table 1). Again, density for ADP could be modelled in one of the three the catalytic sites of the complex (Fig. 2D, *green density*).

**Fig 2.**
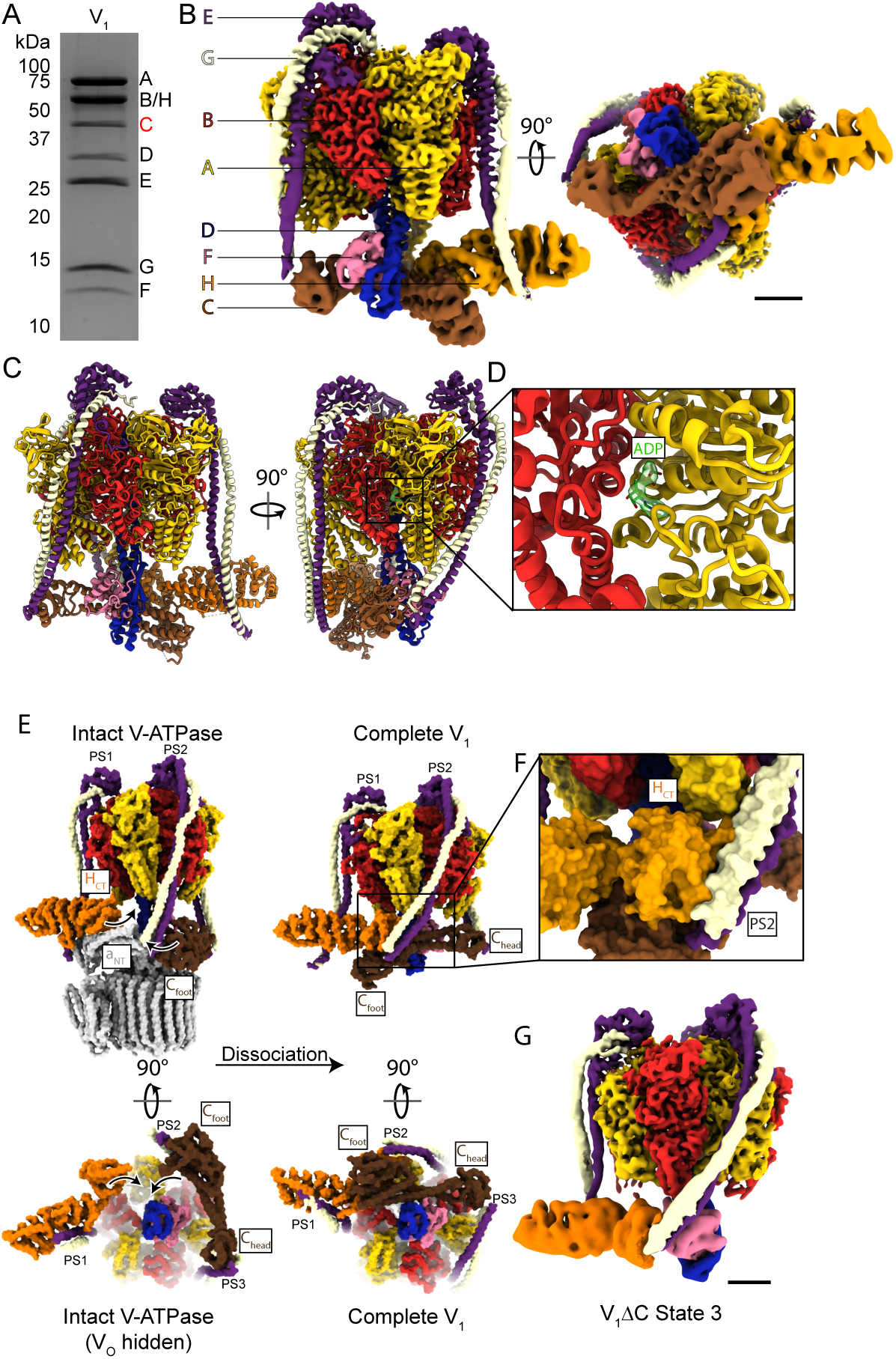
Structure of the complete V_1_ complex. **A**, SDS-PAGE of the purified complete V_1_ complex including subunit C (*red*). **B**, CryoEM map of the complete V_1_ complex. Scale bar, 25 Å. **C**, Atomic model for the complete V_1_ complex shows the complex in rotational State 2. **D**, Density for ADP (*green*) is found in one of the catalytic AB pairs. **E**, Comparison of atomic models for intact V-ATPase (*left*) and complete V_1_ (*right*) shows the conformational changes that occur upon separation of V_1_ from V_O_. **F**, The N terminus of peripheral stalk 2 (PS2) interacts with the C-terminal domain of subunit H (H_CT_) in the complete V_1_ structure. **G**, A subset of the particle images from the complete V_1_ dataset show the V_1_ΔC complex in rotational State 3. Scale bar, 25 Å.

The dissociated but complete V_1_ complex is in rotational State 2, making it incompatible for assembly with the dissociated V_O_ complex in State 3^10,11^. In addition to this mismatch of rotational states, dissociation of V_1_ from V_O_ is accompanied by bending of peripheral stalk 1 (PS1) to bring H_CT_ into contact with peripheral stalk 2 (PS2), and bending of peripheral stalk 3 (PS3) to bring subunit C into contact with the central rotor subunits D and F (Fig. 2E, Supplementary Video 2). This conformation of subunits H and C resembles low-resolution cryoEM of the *Manduca sexta* V_1_ complex^40^, with the C and H interaction similar to a low-resolution cryoEM reconstruction of a yeast CE_3_G_3_H subcomplex^21^. In intact V-ATPase, PS2 is between the N-terminal domain of subunit a (a_NT_) and C_foot_ (Fig. 2E, *top left*). However, in the dissociated complete V_1_ complex, PS2 breaks its interactions with a_NT_ and C_foot_ and is pulled across a neighboring A subunit to form the new interaction with an *α*-helix (residues 460 to 477) in H_CT_ (Fig. 2F). This surface of H_CT_ interacts with a_NT_ in intact V-ATPase and only becomes exposed and available to interact with PS2 after V_1_ separates from V_O_ and H_CT_ rotates toward PS2. Interestingly, a similar H_CT_:PS2 interaction occurs between H_CT_ from one V_1_ΔC complex and PS2 from a symmetry related V_1_ΔC complex in the previously described crystals of V_1_ΔC^27^. As subunit C rotates toward the central stalk upon dissociation of V_1_ from V_O_, C_foot_ moves beneath subunit H into the conformation that brings it into contact with subunit F. The interaction of subunit C with subunit F may contribute to inhibiting ATP hydrolysis in complete V_1_ by preventing rotation of the rotor subcomplex (Fig. 2E, bottom right).

As expected, a subset of the particle images from the complete V_1_ dataset were found without subunit C. These images allowed for calculation of a map of V_1_ΔC at a nominal overall resolution of 4.0 Å, with local resolutions ranging from 3.5 to 9.3 Å resolution (Fig. 2G, Fig. S2B and D).

### The V_1_ ΔC complex is found in all three rotational states

The conformation of V_1_ΔC seen from cryoEM is different from the conformation found in the earlier 6.2 to 6.5 Å resolution crystal structure of the V_1_ΔC complex^27^ in several ways. Most notably, the cryoEM map is in rotational State 3 whereas the crystal structure shows V_1_ΔC in rotational State 2. Further, the conformation of subunit H differs between the cryoEM and crystal structures. In the cryoEM structure, the conformation of subunit H in V_1_ΔC is similar to its conformation in intact V-ATPase^9,25^, while the crystal structure shows a ∼150° rotation of H_CT_ that brings it into contact with an AB pair. Whereas the cryoEM map came from a yeast strain that possessed all the genes for V-ATPase subunits, the crystal structure for V_1_ΔC was obtained using a yeast strain in which the *VMA5* gene that encodes subunit C was deleted. To determine whether the difference in rotational state and subunit H conformation observed in cryoEM was due to subunit C dissociating from the complete V_1_ complex, rather than being absent due to deletion of the subunit C gene, we generated a yeast strain with *VMA5* deleted and a 3×FLAG tag at the C terminus of subunit A (Vma1p) to facilitate V_1_ΔC purification (Fig. 3A). As expected^23^, both the complete V_1_ preparation (Fig. 3B, *purple triangles*) and V_1_ΔC preparation (Fig. 3B, *black diamonds*) lack detectable magnesium ATPase activity.

**Fig. 3.**
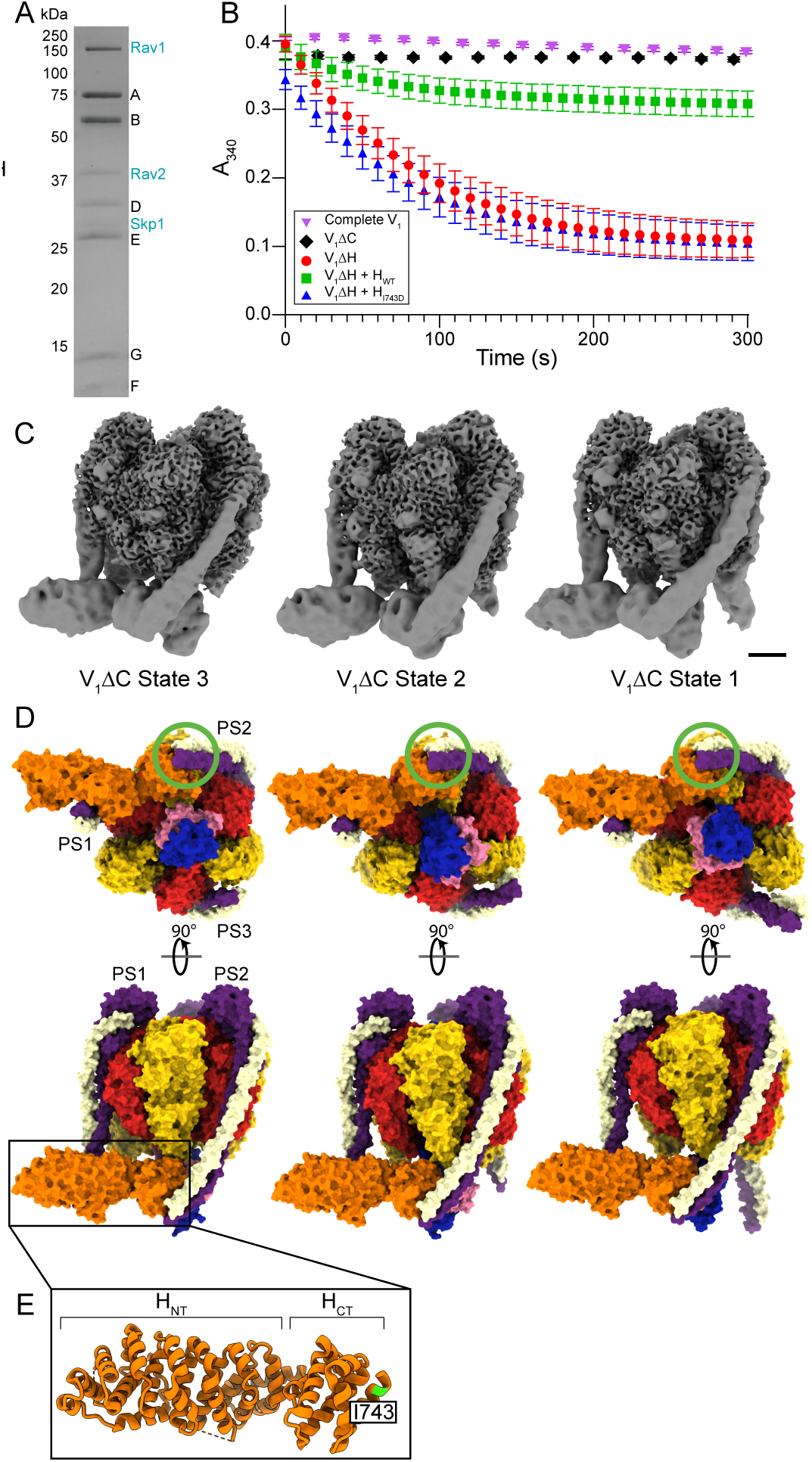
Structures of the V_1_ΔC complex in three rotational states. **A**, SDS-PAGE of V_1_ΔC purified from yeast grown in acidic conditions shows co-purification of the RAVE complex (*RAVE subunits labelled in blue*). **B**, As previously demonstrated, activity assays for complete V_1_ (*purple triangles*) and V_1_ΔC (*black diamonds*) show negligible activity (±s.d., n=3 assays). In contrast, V_1_ΔH (*red circles*) has strong ATPase activity with a 5-fold excess of recombinant wildtype subunit H (*green squares*) inhibiting activity but a 5-fold excess of recombinant subunit H bearing the mutation I743D (*blue triangles*) unable to inhibit (±s.d., n=6 from two separate batches of protein). **C**, CryoEM maps for V_1_ΔC in State 3 (*left*), 2 (*middle*), and 1 (*right*). Scale bar, 25 Å. **D**, Atomic models for V_1_ΔC in State 3 (*left*), 2 (*middle*), and 1 (*right*). Peripheral stalks 1 (PS1), 2 (PS2), and 3 (PS3) are indicated and the interaction between PS2 and the H_CT_ is circled in green. **E**, H_CT_ also interacts with PS2 in the V_1_ cryoEM structures. The location of Ile 743, which prevents ATPase inhibition by subunit H when mutated to Glu, is indicated.

CryoEM of the V_1_ΔC protein preparation (Fig. S3A, Table 1) revealed two main populations of images, corresponding to V_1_ΔC in rotational State 3 (Fig. 3C, *left*), as seen in the V_1_ΔC map calculated from the complete V_1_ dataset, and V_1_ΔC in rotational State 2 (Fig. 3C, *middle*). From the final data set of particle images, 45% contributed to the State 2 map while 55% contributed to the State 3 map. In the presence of 1 mM magnesium ATP (Fig. S3B) V_1_ΔC was found in all three rotation states, including State 1 (Fig. 3C, *right*). From this dataset, 24% of particle images contributed to the State 1 map, 25% to the State 2 map, and 51% to the State 3 map. The identification of all three rotational states for V_1_ΔC suggests that although the complex lacks the magnesium ATPase activity of V_1_ΔH^23^, it is still capable of slowly interconverting between rotary conformations. The fraction of particle images contributing to each rotational state suggests that the complex slightly favours State 3, the conformation that matches the dissociated V_O_ complex.

The cryoEM maps of V_1_ΔC had nominal overall resolutions of 3.3 Å for States 2 and 3 in the absence of ATP, and 3.3, 3.3, and 3.2 Å for rotational States 1, 2, and 3, respectively, in the presence of 1 mM ATP (Fig. S3A and B), with local resolutions of 3.2 to 9.0 Å (Fig. S3C and D). Again, the best resolved regions in the maps were within in the complex’s catalytic core and the poorest resolution was in subunit H and F, with the lower resolution and three-dimensional variability analysis (Supplementary Video 3) suggesting that these subunits remain flexible. These maps enabled construction of atomic models for all three rotational states of V_1_ΔC by a combination of *de novo* model building and flexible fitting of the subunit H and F crystal structures (Fig. 3D, Table 1).

In addition to these three conformations of V_1_ΔC, we found that when sample that had been left at 4°C was imaged, another structure appeared during 3D classification. This additional structure appeared with increasing abundance the longer the sample was left at 4°C, being rare with freshly purified V_1_ΔC (<10% of particles images) but accounting for ∼40% of particle images after leaving the specimen at 4°C for four days. The additional structure consisted of disrupted complexes attached together (Fig. S4A). The cryoEM map of the additional structure fits the crystal structure of V_1_ΔC (Fig. S4B, *yellow*) along with an additional subunit H and subunit EG dimer from a contacting V_1_ΔC in the crystal lattice^27^ (Fig. S4B, *blue*). Thus, it appears that the subunit H conformation and interactions seen in crystals of V_1_ΔC, which took approximately two weeks to grow^27^, gradually occur in solution, explaining the difference in subunit H conformation seen by cryoEM and X-ray crystallography. In addition to the V_1_ΔC structure seen by crystallography, the cryoEM map of the additional structure also contains a third subunit H, which matches the conformation of subunit H in cryoEM of freshly purified V_1_ΔC (Fig. S4C, *red*).

The finding that V_1_ΔC can adopt all three rotational states shows that removal of subunit C allows the complex to interconvert between the states, but not at a rate sufficient to allow rapid ATP hydrolysis. In contrast, subunit H is required for inhibition of ATP hydrolysis in V_123_, with purified V_1_ΔH^25^ showing strong but non-linear ATPase activity^23,25^ (Fig. 3B, *red circles*). The proximity of H_CT_ to subunit F when V_1_ΔC is in rotational State 1 or 3 explains how these subunits can be crosslinked in V_1_ but not in intact V-ATPase^41^. However, the absence of interaction between H_CT_ and subunit F in State 2, even in the presence of ATP, and 3D variability analysis that shows movement of H_CT_ relative to subunit F in States 1 and 3 (Supplementary Video 3), suggests that interaction of H_CT_ with subunit F is not responsible for inhibiting ATP hydrolysis in V_1_ΔC. In comparison, H_CT_ forms an interaction with PS2 in the complete V_1_ structure (Fig. 2F) and all three rotational states of the V_1_ΔC complex (Fig. 3D, *green circles*). This H_CT_:PS2 interaction distorts the complex, pulling PS2 across a catalytic A subunit (Fig. 2E, *right*, and 3D, *top*). Therefore, we considered the possibility that interaction of H_CT_ with PS2 inhibits ATP hydrolysis by preventing catalytic conformational changes, reminiscent of the partial inhibition of V-ATPase by the bacterial effector protein SidK^42^. To test this hypothesis, we expressed and purified recombinant subunit H, both as wildtype and bearing the point mutation I473D, which introduces a negative charge into the hydrophobic patch on the surface of subunit H where it interacts with PS2 (Fig. 3E). As described previously^23,25^, recombinant subunit H incubated with V_1_ΔH inhibits the complex’s ATPase activity^25^ (Fig. 3B, *green squares*). However, the H construct bearing the I473D mutation showed no ability to inhibit ATP hydrolysis in V_1_ΔH (Fig. 3B, *blue triangles*), supporting the hypothesis that interaction of H_CT_ with PS2 inhibits ATP hydrolysis.

### The RAVE complex co-purifies with V_1_ΔC

As seen previously^34^, SDS-PAGE of the V_1_ΔC preparation showed additional bands on the gel that corresponded in molecular weight to the subunits of the RAVE complex (Fig. 3A, *blue*). We found that these bands were particularly enriched when yeast were cultured in acidic conditions, known to favour growth with a defective or missing gene for a V-ATPase subunit^43^. Mass spectrometry (Table S2) confirmed the identity of the bands as Rav1p, Rav2p, and Skp1p. However, despite significant effort, we were unable to obtain a cryoEM map of V_1_ with RAVE bound. The binding of RAVE to V_1_ΔC suggests a frustrated RAVE complex that is unable to assemble V_1_ with V_O_. Consistent with this hypothesis, RAVE copurifies with V_1_ΔC most noticeably when yeast are cultured in acidic conditions where the effect of subunit C deletion on yeast growth is minimized (Fig. 3A). Binding of subunit C to the RAVE-V_1_ complex is also known to be tighter than binding of subunit C to the RAVE complex alone *in vitro*^34^. Therefore, as previously suggested^33,34^, these findings suggest that RAVE binds V_1_ΔC and recruits subunit C to reassemble the intact V-ATPase.

## Discussion

The structures described above suggest a sequence of events that occurs during the regulated separation and reassembly of the V_1_ and V_O_ complexes (Supplementary Video 4). During ATP hydrolysis and proton pumping, intact V-ATPase cycles sequentially between the three observed rotational States 1, 2, and 3 (Fig. 4A). Upon glucose depletion the V_1_ and V_O_ regions separate^13^. As this separation results in the V_O_ complex adopting State 3^10^ and the complete V_1_ complex adopting State 2 (Fig. 2), it is not clear whether separation of the intact V-ATPase occurs from State 2 or from State 3. If separation occurs when V-ATPase is in State 2, the V_O_ region would have to undergo a conformational change to adopt the experimentally observed State 3. This transition could occur due to rotation of V_O_ ‘backwards’ to its autoinhibited state, from State 2 → State 1 → State 3, driven by the proton motive force established previously during proton pumping. Alternatively, separation of V_1_ from V_O_ could occur when intact V-ATPase is in State 3 (modelled in Fig. 4B). In this case, the complete V_1_ complex would have to transition to State 2 after separation, likely driven by ATP hydrolysis from State 3 → State 1→State 2 before it adopts its observed conformation. Through either route, the separation of V_1_ from V_O_ leads to mismatched conformations, with the V_O_ subcomplex in State 3 and the complete V_1_ complex in State 2 (Fig. 4C). This rotational mismatch, as well as the dramatic change in conformation of subunit H, C, and the peripheral stalks of complete V_1_, prevent spontaneous reassembly of V_1_ and V_O_.

**Fig. 4.**
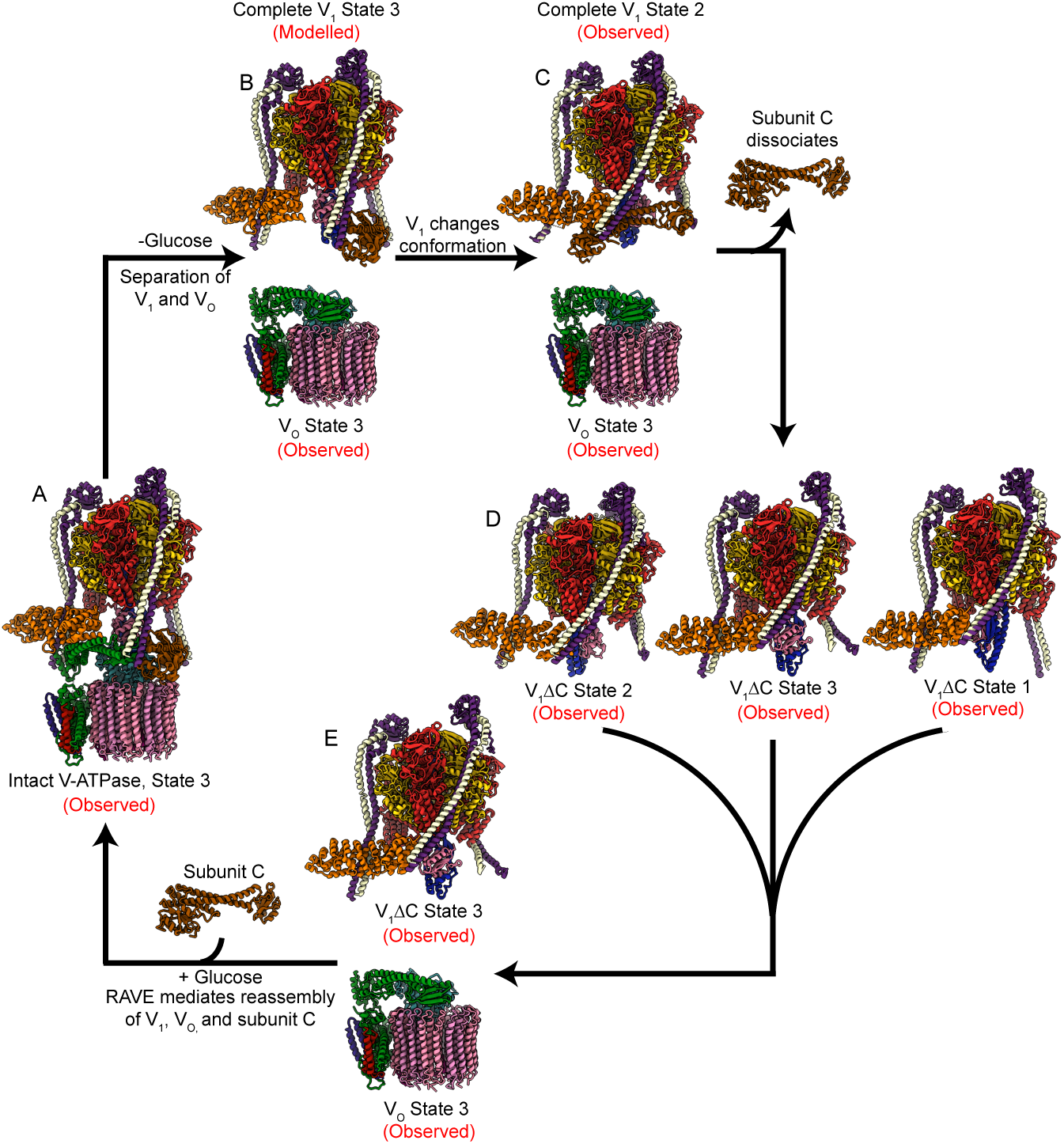
Cycle of conformational changes in V-ATPase reversible dissociation. **A**, Intact active V-ATPase cycles between rotational States 1, 2, and 3 during ATP hydrolysis and proton pumping. **B**, Upon glucose depletion the V_1_ and V_O_ regions separate. **C**, The complete V_1_ complex transitions to State 2 and adopts an inhibited conformation, leading to a mismatch of rotational states for V_1_ and V_O_, which prevents premature reassembly. **D**, Following dissociation of subunit C, V_1_ΔC can sample all three rotational states. V_1_ΔC in State 3 is competent for reassembly with the V_O_ complex in State 3, but the absence of subunit C prevents spontaneous reassembly. **E**, The RAVE complex reassembles V_1_ΔC in State 3 with V_O_ in State 3 and subunit C.

The structure of the complete V_1_ in State 2 suggests that the complex cannot adopt State 3 without dissociation of subunit C. Once subunit C dissociates, V_1_ΔC can sample all three rotational states (Fig. 4D), albeit without the rapid interconversion that leads to high rates of ATP hydrolysis in V_1_ΔH or intact V-ATPase. V_1_ΔC complexes in State 3 (Fig. 4E) are competent for binding to the V_O_ complex in State 3, but the absence of subunit C prevents spontaneous reassembly. Recruitment of subunit C to V_1_ΔC requires the RAVE complex, as described previously^34^ and as illustrated by the co-purification of RAVE with V_1_ΔC complexes in conditions that favor yeast growth (Fig. 3A). The RAVE complex reassembles V_1_ΔC in State 3 with subunit C and V_O_ in State 3 to form a functional intact V-ATPase (Fig. 4A).

The reversible dissociation of V_1_ and V_O_ is remarkable in that the connection between the two complexes must be mechanically robust during ATP-hydrolysis-driven proton pumping, but also sufficiently labile to permit disruption for regulation of V-ATPase activity. The structures described here illustrate how these conflicting requirements are met through a coordinate series of conformational changes in the V_1_ and V_O_ complexes, including dissociation and RAVE-mediated recruitment of subunit C to V_1_. This ‘on’ or ‘off’ control of V-ATPase activity represents one level of regulation of the enzyme activity in response to changing conditions, with finetuning of activity achieved through incorporation of different subunit isoforms and interactions with specific lipids or other molecules^44–46^.

## Acknowledgements

We thank Samir Benlekbir, Yong Zi Tan, and Hui Guo for assistance with cryoEM data collection. We thank Patricia Kane for the gift of the PMK549 yeast strain and for discussions about yeast growth, and Stephan Wilkens and Rebecca Oot for discussions about the V_1_ΔC crystal structure. TV was supported by an Ontario Graduate Scholarship, KAK was supported by an Ontario Graduate Scholarship and a Canada Graduate Scholarship, and JLR was supported by the Canada Research Chairs program. This work was supported by Canadian Institutes of Health Research grant PJT166152 (JLR). CryoEM data was collected at the Toronto High-Resolution High-Throughput cryo-EM facility, supported by the Canada Foundation for Innovation and Ontario Research Fund. Mass spectrometry data was collected at the Network Biology Collaborative Center at the Lunenfeld-Tanenbaum Research Institute.

## Author contributions

TV purified V_1_, V_1_ΔC, V_1_ΔH, and recombinant subunit H constructs, and performed enzyme assays, cryoEM, and the associated analysis with those specimens. KAK purified intact V-ATPase and performed the cryoEM and associated analysis for that specimen. SAB prepared the *VMA1*-3×FLAG, *vma5Δ* yeast strain. MCJ prepared protein for early experiments and advised on protein purification. JLR supervised and coordinated experiments

TV and JLR wrote the manuscript and prepared figures with input from the other authors.

## Materials and Methods

### Construction of yeast strain for V_1_ΔC purification

The yeast strain SABY119 (VMA1 3×FLAG, *vma5Δ*), was generated by homologous recombination to insert a 3×FLAG tag C-terminal to the *VMA1* gene in the background strain PMK549 (*vma5Δ::kanMX* in BY4741), which was a gift from Prof. Patricia Kane (SUNY Upstate Medical University). DNA sequence for the 3×FLAG tag followed by a *URA3* marker was amplified from plasmid pJT1 with primers 5′-TCG AAA AAT TGT TGA GCA CTA TGC AAG AAA GAT TTG CTG AAT CTA CCG ATG ACT ACA AAG ACC ATG ACG G-3′ and 5′-GAA GAA AAG ACA TCT AAC AAA TAT ACC AGA AGA TAA ATG CTA CAT ATA TCA TAT CAT CGA TGA ATT CGA GCT CG-3′ and the PCR product was used to transform PMK549 as described previously^25^. Transformed cells were selected by growth on minimal medium lacking uracil and successful transformants were identified by PCR.

### Construction of plasmid and purification of subunit H

The plasmid pTV27 was generated to introduce the point mutation I473D in the *VMA13* (subunit H) gene on plasmid pSAB13, which was designed for expression of wildtype subunit H^25^. The primers 5′-GCA GGC AAT CGA TGG ATA TAC CTT CAA ATA AGG ATC CG-3′ and 5′-CCG CCT TTC TCC CTT CGG GAA GCG TG-3′; and 5′-AAG GGA GAA AGG CGG ACA GGT ATC CG-3′ and 5′-TAT CCA TCG ATT GCC TGC GTG GCC TTG-3′ were used to generate Gibson assembly fragments from pSAB13, which were assembled using the Gibson Assembly Cloning Kit (New England Biolabs). Recombinant H_WT_ and H_I473D_ were expressed in *E. coli* and purified as described previously^25^.

### Purification of intact V-ATPase

Yeast strain SABY31 (VMA1-3×FLAG, *stv1Δ*)^25^ in YPD media (11 L) was grown in a Microferm fermenter (New Brunswick Scientific) for 36 hours (OD_660_=6 to 9) at 30°C with stirring at 300 rpm and aeration at 34 cubic feet per hour. Thirty min prior to harvest, 22% (w/v) D-glucose solution (1 L) was added to promote assembly of V-ATPase. All subsequent steps were performed at 4 °C. Cells were lysed and membranes were harvested and solubilized with n-dodecyl β-D-maltopyranoside (DDM) as described previously^25^. Intact V-ATPase was isolated from solubilized membranes by affinity purification, using a gravity column containing 0.8 mL of anti-FLAG M2 Affinity gel (Sigma-Aldrich) equilibrated with DTBS (50 mM Tris-HCl pH 7.4, 150 mM NaCl, 0.05% [w/v] DDM). The column was washed with ten bed volumes of High-LTBS buffer (50 mM Tris-HCl pH 7.4, 150 mM NaCl, 0.01% [w/v] lauryl maltose neopentyl glycol [LMNG]) and one bed volume of Low-LTBS buffer (50 mM Tris-HCl pH 7.4, 150 mM NaCl, 0.002% [w/v] LMNG). V-ATPase was eluted with 1.5 mL of Low-LTBS buffer containing 150 µg/mL 3×FLAG peptide and 0.5 mL of Low-LTBS buffer without 3×FLAG peptide. Purified V-ATPase in buffer containing LMNG was concentrated to 1.9 mg/mL with a 100 kDa MWCO VivaSpin 6 centrifugal concentrator at 2000 ×g for 15 min (Sartorius), followed by a VivaSpin 500 concentrator at 12,000 ×g for 2 min.

### Purification of complete V_1_ and V_1_ΔC

For purification of complete V_1_, two 1 L cultures of the yeast strain SABY31 in YPD were grown for 40 h at 30°C in Fernbach flasks with shaking at 220 rpm (OD_660_=6 to 7). For purification of V_1_ΔH, the yeast strain SABY42^25^ was grown as described above. For purification of V_1_ΔC, the yeast strain SABY119 was grown as described above but with shaking at 180 rpm and in YPD media adjusted to pH 5 by addition of HCl. All subsequent steps were performed at 4°C. Cells were harvested by centrifugation at 4000 ×g for 15 min with an Avanti J-15R Benchtop centrifuge and resuspended in lysis buffer (phosphate-buffered saline pH 7.4, 8% [w/v] sucrose, 2% [w/v] sorbitol, 2% [w/v] glucose, 5 mM 6-aminocaproic acid, 5 mM benzamidine hydrochloride, 5 mM EDTA, and 0.001% [w/v] PMSF) at 2 mL/g cell pellet. Cells were lysed with an Avestin Emulsiflex C3 Homogenizer at 20,000 to 25,000 psi for 9 rounds, resting on ice for 5 min between each round. Cellular debris was pelleted by centrifugation at 3000 ×g for 10 min and membranes removed by ultracentrifugation at 139,000 ×g or 40 min. The supernatant was filtered with a 0.45 µm syringe filter, divided into thirds, and applied to three gravity columns each containing 0.8 mL of anti-FLAG M2 Affinity gel and equilibrated with TBS (50 mM Tris-HCl pH 7.4, 150 mM NaCl). The columns were washed with ten bed volumes of TBS and the sample was eluted from the column with three bed volumes of TBS containing 3×FLAG peptide at 150 µg/mL and a final addition of 1 bed volume of TBS. The sample was concentrated to a final volume of 300 µL with a 100-kDa MWCO Vivaspin 6 centrifugal concentrator. Samples for cryoEM were further purified with a Superose 6 Increase 10/300 GL column. Fractions were assessed by SDS-PAGE and those containing V_1_ were pooled and concentrated to ∼5.7 mg/mL for complete V_1_ and 3 to 4 mg/mL for V_1_ΔC before analysis by cryoEM.

### CryoEM specimen preparation

All samples were applied to homemade holey gold grids with regular arrays of holes^47^. For intact V-ATPase, grids were glow discharged in air for 2 min and 3 µL of sample was applied in a modified FEI Vitrobot at ∼100% relative humidity and 4°C. Grids were blotted for 11 s and plunge frozen in a mixture of liquid ethane and liquid propane^48^. For complete V_1_, IGEPAL (Sigma) was added to a final concentration of 0.025% (v/v) prior to freezing and 2 µL of sample was applied to the grid. Grids were blotted for 5 s with the Vitrobot and plunge frozen after waiting for 1 s. For V_1_ΔC grid preparation, IGEPAL was added to a final concentration of 0.025% and 1.5 µL of sample was applied to grids that were glow discharged in air for 15 s. These grids were blotted for 2 s with a Leica EM GP2 grid freezing device and plunged in liquid ethane. When used, MgATP was added to a final concentration of 1 mM immediately prior to sample application for each grid.

### Data collection

All samples were screened with an FEI Tecnai F20 electron microscope operating at 200 kV and equipped with a Gatan K2 Summit direct detector device camera. Images were collected with K2 Summit operating in counting mode at 25,000× magnification, corresponding to a pixel size of 1.45 Å. Movies were recorded as 30 frames over 15 s, with an exposure rate of ∼5 electrons/pixel/s. High-resolution data collection was performed on a Titan Krios G3 electron microscope (Thermo Fisher) operating at 300 kV and equipped with a Falcon 4 camera. Movies were collected with 30 exposure fractions at a nominal magnification of 75,000×, corresponding to a calibrated pixel size of 1.03 Å. Further details for each dataset are given in Table 1. Automated data collection was performed with the *EPU* software package.

### Image analysis

Movie alignment and estimation of CTF parameters in patches was performed on-the-fly with *cryoSPARC Live*^49^. Templates for particle selection were generated by 2D classification of manually selected particle images. Non-stringent settings for image power and correlation were used for template-based particle picking to maximize the number of particle views selected and minimize the loss of rare views of the complex. Individual particle motion correction was performed^50^ and datasets were cleaned with multiple rounds of 2D classification and ab initio 3D classification in *cryoSPARC v2*^37^.This process provided 384,031 particle images for intact the V-ATPase dataset, 315,308 particle images for the complete V_1_ dataset, 511,582 particle images for the V_1_ΔC dataset, and 418,164 particle images for the V_1_ΔC with 1 mM MgATP dataset.

For intact V-ATPase, ab initio 3D classification and heterogeneous refinement led to the identification of classes corresponding to the three main rotational states of the complex. These classes were refined with nonuniform refinement^51^ followed by CTF refinement and masked local refinement of the V_1_ region. Local refinement of the V_O_ region was performed following signal subtraction to remove the rest of the complex from images. For the complete V_1_ complex, ab initio 3D classification and heterogeneous refinement led to the identification of the complete V_1_ complex in State 2 and V_1_ΔC in State 3, which were refined with nonuniform refinement. For V_1_ΔC, ab initio structure determination and 3D classification with heterogeneous refinement allowed identification of V_1_ΔC in State 3, V_1_ΔC in State 2, and the aggregated and disrupted V_1_ΔC complex (Fig. S4), which were refined with nonuniform refinement. When 1 mM MgATP was added to the V_1_ΔC sample, an additional class showing the complex in State 1 was found and refined with nonuniform refinement.

### Construction of atomic models

Atomic models were constructed by manual model building in *Coot*^52^ followed by several rounds of real space refinement with *PHENIX*^53^ and refinement using *ISOLDE*^54^. For intact V-ATPase, previously published models 3J9T^9^ and 6M0R^11^ were used to begin modelling. The models for States 2 and 3 were built by rigid body fitting of the model for State 1 into the maps with *UCSF Chimera*^55^, followed by manual model building and refinement as described above. For the V_1_ structures, the previously published model 3J9T^9^ was used as a starting model, with the crystal structure for subunit H and C from 1HO8^26^ and 1U7L^56^, respectively. Figures were rendered with *Chimera*^55^ and *UCSF ChimeraX*^57^.

### Activity assays

Enzyme-coupled ATPase activity assays were performed as described previously^46,58^ with some modifications. Assays were performed in a 96-well plate with a total reaction volume of 160 µL. Purified V_1_ was added to the ATPase assay reaction buffer (50 mM Tris pH 7.4, 3 mM MgCl, 0.2 mM NADH disodium salt, 3.2 units pyruvate kinase, 8 units L-lactic dehydrogenase) and the reaction was initiated with the addition of 1 mM ATP disodium salt and 1 mM phosphoenol pyruvic acid monopotassium salt. When used, recombinant subunit H_WT_ or H_I473D_ were added at 5-fold molar excess to V_1_. Absorbance at 340 nm was monitored at 30°C to measure the signal from NADH.

### Mass spectrometry of RAVE subunits

Mass spectrometry experiments were performed at the Network Biology Collaborative Center of the Lunenfeld-Tanenbaum Research Institute (Toronto) with their standard protocols and analysis pipelines. Briefly, bands were cut from a 15% SDS-PAGE gel and digested with trypsin. For data-dependent acquisition (DDA) LC-MS/MS, digested peptides were analyzed with a nano-HPLC coupled to a mass spectrometer. Data generated were stored, searched, and analyzed with the *ProHits* laboratory information management system platform. The data were then searched using *Mascot* (V2.3.02) and *Comet* (V2016.01 rev.2). The spectra were searched with the *SwissProt* database from May 2012 for a total of 536,029 entries.

## Figures

**Supplementary Fig 1.**
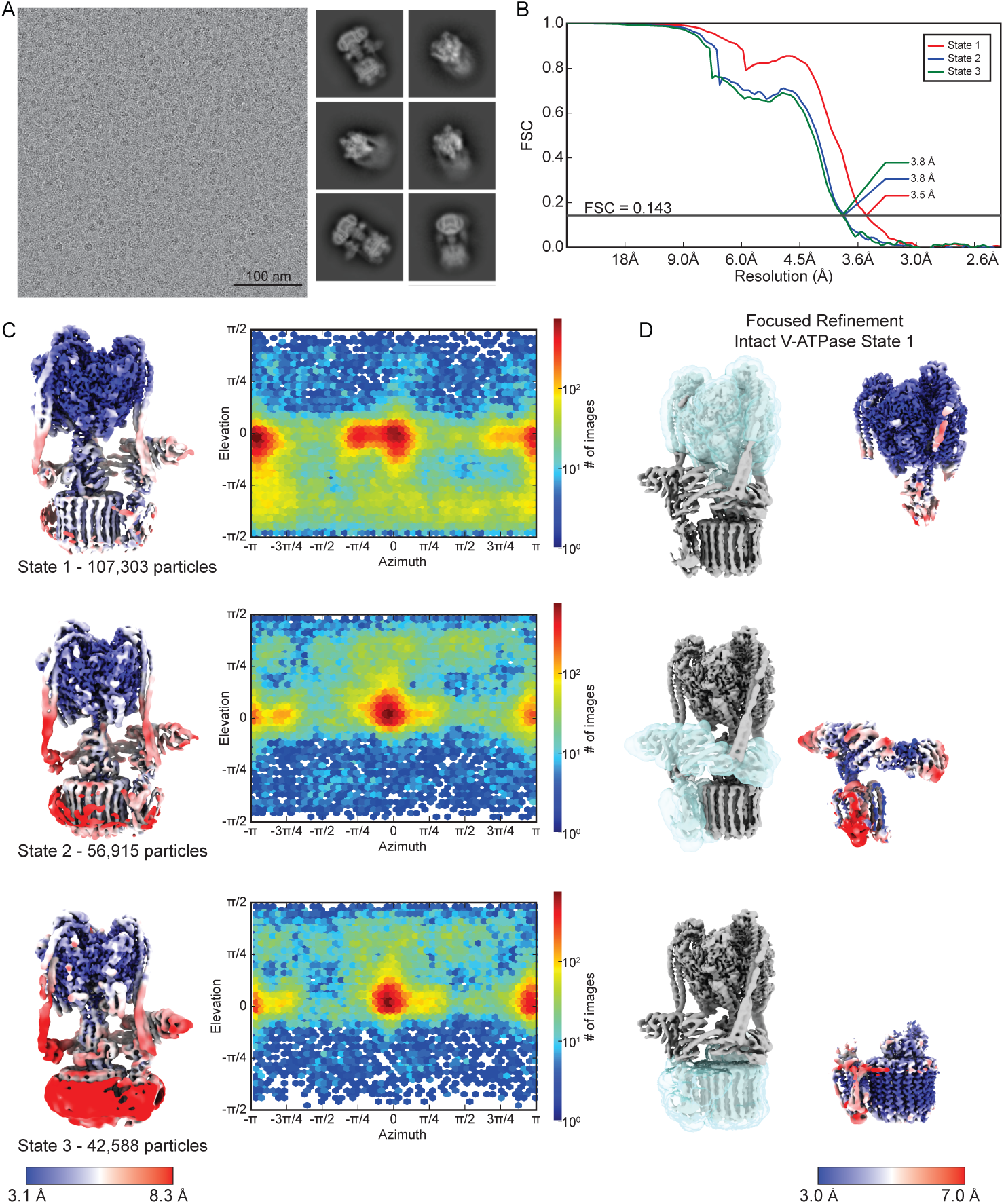
CryoEM of intact V-ATPase. **A**, Example micrograph and 2D class average images. **B**, Fourier shell correlation curves after gold-standard refinement and correction for masking for the three rotational states of intact V-ATPase. **C**, Local resolution maps and orientation distribution plots for rotational State 1 (*top*), State 2 (*middle*), and State 3 (*bottom*). **D**, Masks (*light blue density*) used for local refinement of the V_1_ region (*top*), subunit a and the collar region (*middle*), and the V_O_ region (*bottom*) for V-ATPase (shown for State 1).

**Supplementary Fig 2.**
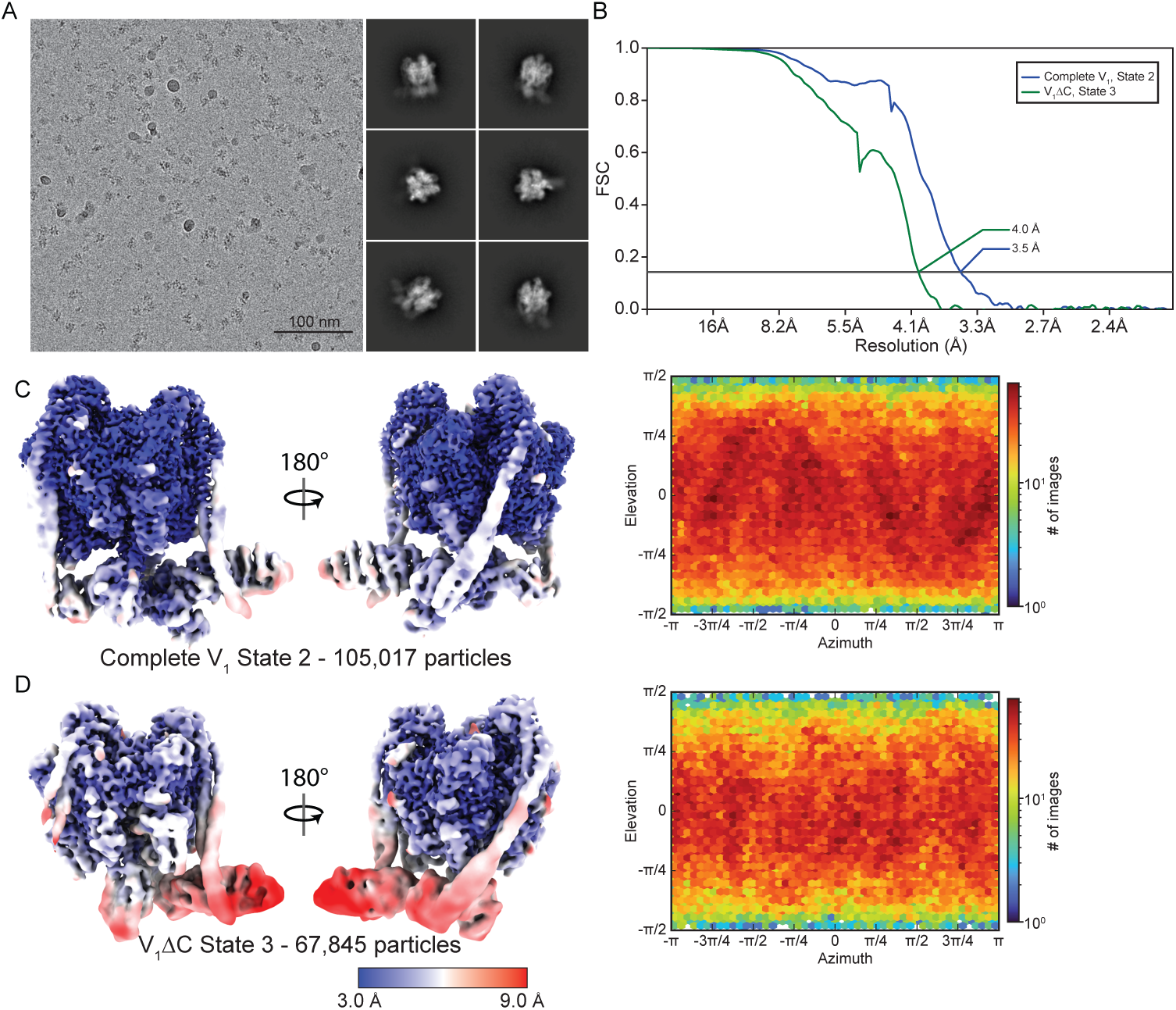
CryoEM of the complete V_1_ complex. **A**, Example micrograph and 2D class average images. **B**, Fourier shell correlation curves after gold-standard refinement and correction for masking for the complete V_1_ complex and V_1_ΔC from the complete V_1_ protein preparation. **C**, Local resolution maps and orientation distribution plot for complete V_1_ in State 2. **D**, Local resolution maps and orientation distribution plot for V_1_ΔC in State 3.

**Supplementary Fig 3.**
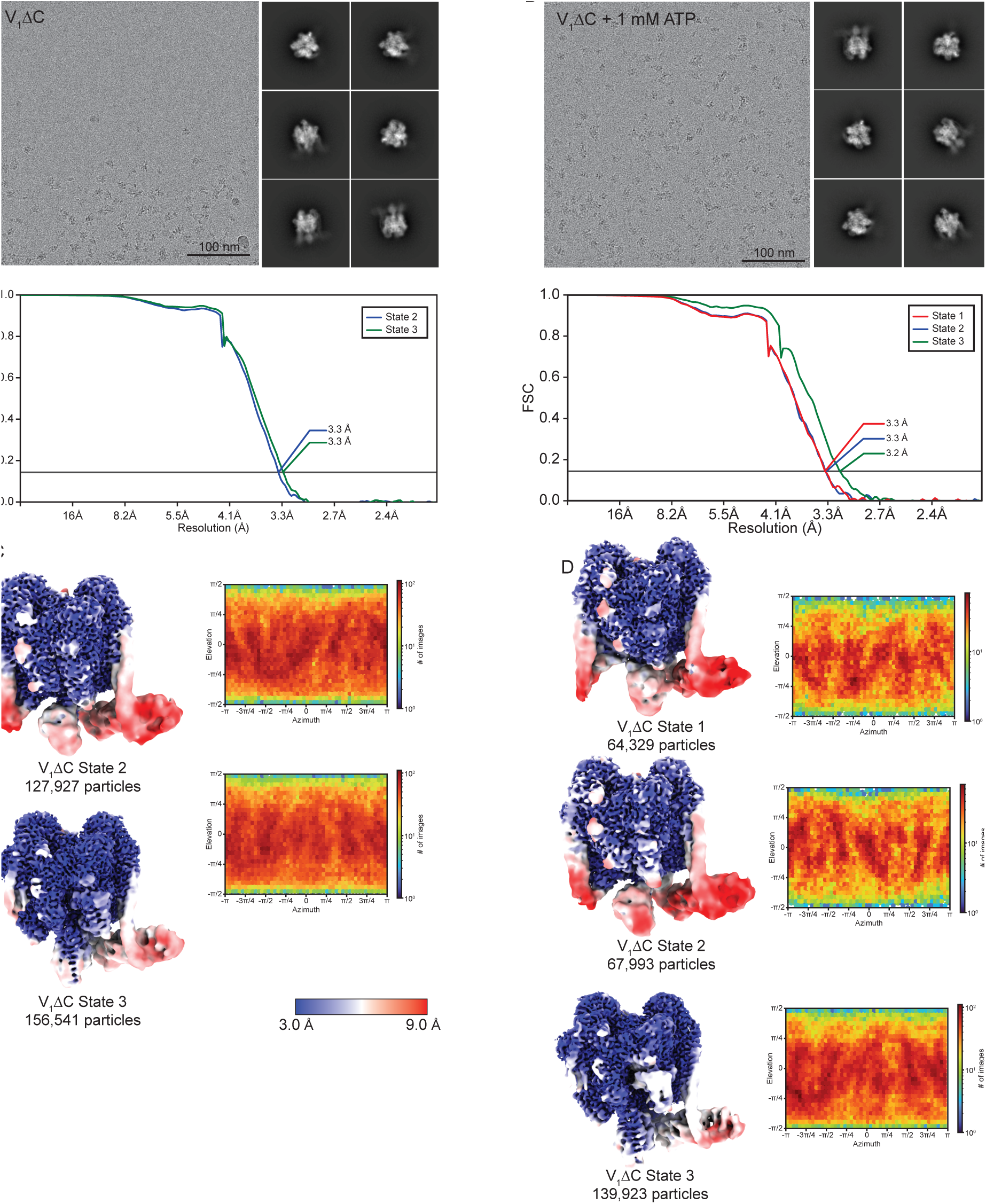
CryoEM of the V_1_ΔC complex. **A**, Example micrographs, 2D class average images, and Fourier shell correlation curves after gold-standard refinement and correction for masking for V_1_ΔC in States 2 and 3. **B**, Example micrograph, 2D class average images, and Fourier shell correlation curves after gold standard refinement and correction for masking for V_1_ΔC with 1 mM ATP in States 1, 2, and 3. **C**, Local resolution maps and orientation distribution plot for V_1_ΔC in State 2 (*top*) and State 3 (*bottom*). **D**, Local resolution maps and orientation distribution plot for V_1_ΔC with 1 mM ATP in State 1 (*top*), State 2 (*middle*), and State 3 (*bottom*).

**Supplementary Fig 4.**
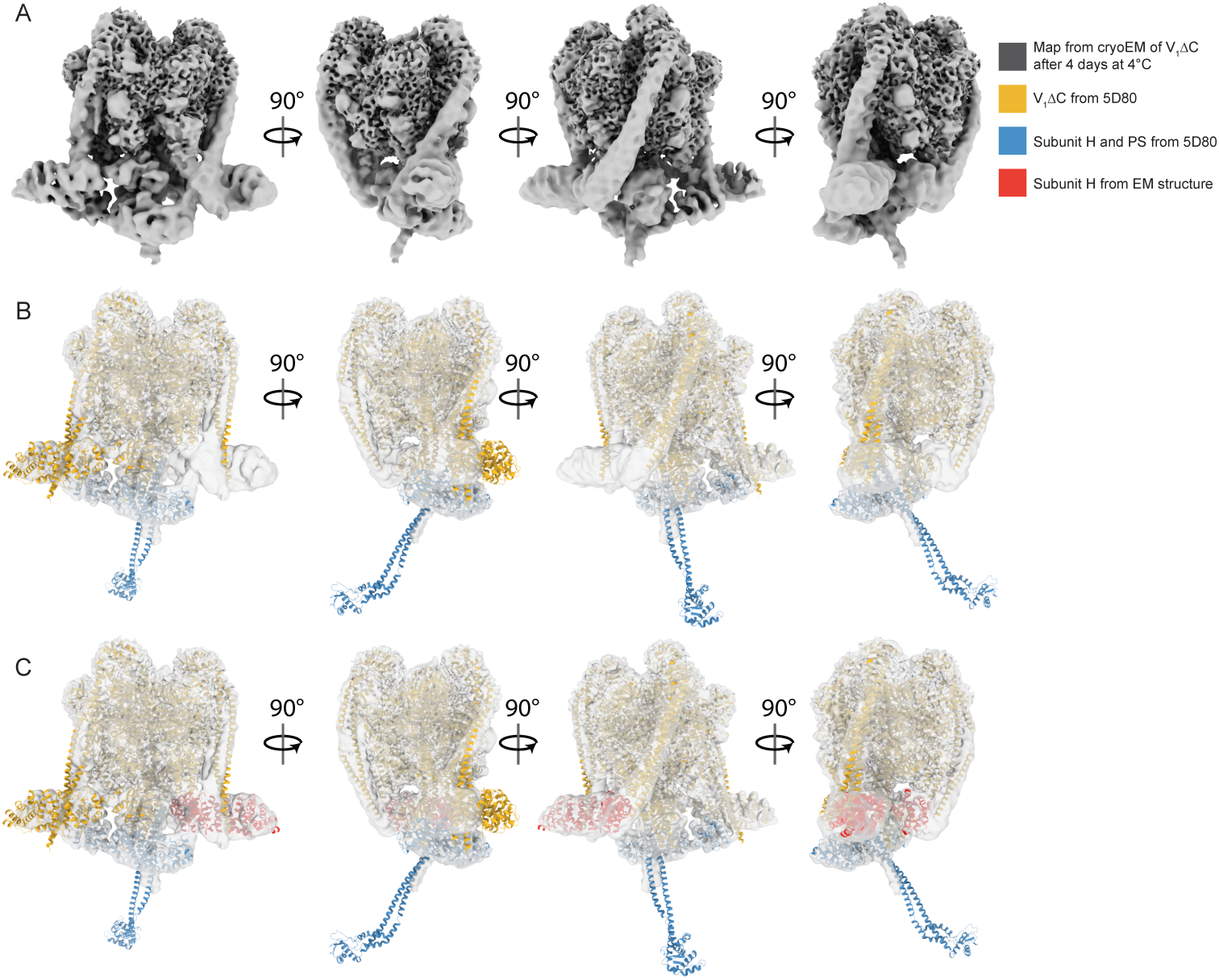
Additional structure of V_1_ΔC complex that appears when the protein preparation is left at 4°C. **A**, CryoEM of the V_1_ΔC sample showed an additional structure that increased in abundance the longer the sample was left at 4°C. **B**, The additional structure fits the model of the V_1_ΔC complex from X-ray crystallography^27^ (*orange*) along with an additional peripheral stalk and subunit H from an adjacent V_1_ΔC complex in the crystal lattice (*blue*). **C**, The entire map of the additional structure is explained by V_1_ΔC from the crystal (*orange*), an adjacent subunit H and peripheral stalk from the crystal (*blue*), and a third subunit H (*red*) not found in the crystal, but matching the conformation of subunit H observed in the cryoEM maps.

**Supplementary Video 1. Three-dimensional variability analysis shows flexibility in the collar region of the complete V**_**1**_ **complex**.

**Supplementary Video 2. Interpolation between the structure of V**_**1**_ **within intact V-ATPase and in the complete V**_**1**_ **complex shows the conformational changes that occur in V**_**1**_ **upon separation from V**_**O**_.

**Supplementary Video 3. Three-dimensional variability analysis shows flexibility in subunit H and the peripheral stalks of the V**_**1**_**ΔC complex in all three rotational states**.

**Supplementary Video 4. Model for the conformational changes that occur during the cycle of dissociation and reassembly of the V**_**1**_ **and V**_**O**_ **complexes**.

